# A developmentally inspired computational model of face recognition that learns continuously through generative memory replay

**DOI:** 10.1101/2025.11.25.690208

**Authors:** Naphtali Abudarham, Galit Yovel

## Abstract

Deep convolutional neural networks (DCNNs) have emerged as powerful models for human face recognition, capturing several hallmark behavioral phenomena such as the face inversion effect and the other-race effect. Yet, key differences remain between how DCNNs and humans, particularly infants, learn to recognize faces. In this study, we present a developmentally-inspired model that addresses three critical gaps between humans and standard DCNNs: (1) DCNNs are trained via one-time batch learning, whereas infants encounter faces gradually over time; (2) DCNNs are trained on thousands of face images with high variability of static images for each identity, while infants are initially exposed to a small number of identities, often limited to close caregivers, through a continuous, dynamic stream of visual input with lower within-identity variability; (3) The goal of DCNNs is to recognize untrained (unfamiliar) faces, while the goal of human face system is to recognize familiar faces. To better approximate developmental face learning, we introduce a model comprising three interacting modules: an Embedder – a DCNN trained to encode face identity continually on up to 10 identities, based on images taken from videos of a TV series; an Autoencoder used for generative memory-replay, to construct images of missing identities in each training phase; and a Memory module – stores identity-specific latent codes. Results show that this model achieves recognition performance comparable to batch-trained DCNNs. We propose this new model as a framework for studying the developmental and mature mechanisms of human face recognition.

## Introduction

Computational models of face recognition offer a framework for exploring the mechanisms that enable humans to achieve this remarkable ability. About a decade ago, deep convolutional neural networks (DCNNs), trained on large-scale datasets of facial images, have reached human-level performance on face recognition tasks. These models were shown to produce internal representations that are aligned with properties of primate visual system (Grossman et al., 2019; Khaligh-Razavi & Kriegeskorte, 2014; Yamins et al., 2014). Moreover, face-trained DCNNs reproduce key behavioral phenomena in human face recognition such as the face inversion effect (Dobs et al., 2023; Yin, 1969; Yovel et al., 2023), the other-race effect (Dobs et al., 2023; O’Toole & Castillo, 2021), sensitivity to critical facial features used by humans for face recognition (Abudarham et al., 2019) and hierarchical identity representation (Abudarham & Yovel, 2021). These parallels have led to a growing interest in using DCNNs as models for understanding the computational principles underlying face recognition in the human mind and brain (for reviews see O’Toole & Castillo, 2021; Phillips & White, 2025; van Dyck & Gruber, 2023)

Yet, important differences remain between DCNNs and humans. Particularly, fundamental differences exist between the training of DCNNs and the learning processes of infants in face recognition. These differences pertain to the nature of learning, the training data set and the goal of the face recognition system (see Table 1). Unlike DCNNs that are trained on large face datasets in a batch mode, all at once, infants acquire facial knowledge continually, under ecological and social constraints. Moreover, studies using head-mounted cameras on infant foreheads during the first year of life, show that infants are typically exposed to a limited set of familiar individuals—primarily caregivers and close family members—encountered repeatedly (Smith & Slone, 2017). These encounters occur in dynamic, real-world settings, where faces are viewed in motion as continuous streams of related frames, naturally varying in pose, illumination, and expression, yet constrained by the consistent background and context of the scene. Finally, the goal of infant face recognition is to recognize the familiar faces of their caregivers and the people they socially interact with, while the goal of DCNN training is to recognize unfamiliar faces – faces that were not included in the training set.

**Table 1:**
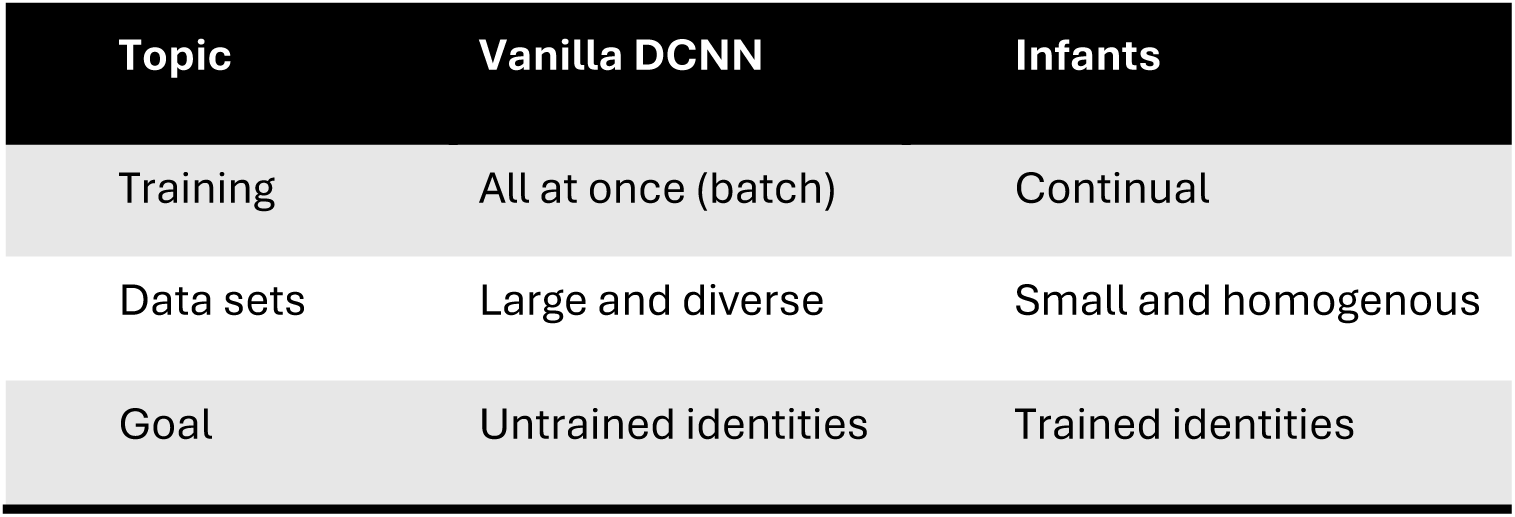
Three main gaps between standard DCNN and infant face learning addressed by the proposed model.

To bring artificial models of face recognition into closer alignment with human development, it is necessary to simulate gradual exposure to different identities. When a vanilla DCNN is trained sequentially like humans, first on one set of identities (e.g., parents) and later, on a different set (e.g., grandparents), without explicit mechanisms to maintain access to the earlier examples, the optimization process adjusts the network’s weights to improve performance on the new inputs. In doing so, the internal representational geometry becomes reorganized to favor the new identities, often at the expense of discriminability for previously learned ones. Consequently, earlier knowledge is effectively lost. This well-known phenomenon is known as catastrophic forgetting (French, 1999; McCloskey & Cohen, 1989).

The **Complementary Learning Systems (CLS)** theory offers a formal framework of how the brain achieves continual learning without catastrophic forgetting (Kumaran et al., 2016; McClelland et al., 1995). According to this theory, the hippocampus supports rapid encoding of individual experiences, while the neocortex gradually abstracts regularities across multiple episodes through slower, integrative learning. Replay of the individual experiences acts as the critical interface between these systems: hippocampal reactivation of recent experiences trains neocortical networks offline, enabling consolidation of new information, without overwriting existing representations. These ideas are supported by animal studies that found that hippocampal place-cell sequences, which encode behavioral trajectories, are reactivated during sharp-wave ripples in both slow-wave sleep and quiet wakefulness, occurring at compressed time scales (Carr et al., 2011). This replay is suggested to support systems-level consolidation by reinforcing weak or vulnerable memories and reducing interference with newly acquired content (Ólafsdóttir et al., 2018). In primates and humans, replay-like reactivations predict later memory performance and appear to prioritize salient or weakly encoded events (Carr et al., 2011; Schapiro et al., 2018). Functional reuse of these replay events in neocortical areas suggests a mechanism for memory integration and stability (Ólafsdóttir et al., 2018).

Drawing on these ideas, memory replay was suggested as a compelling approach to mitigate catastrophic forgetting in continual training of artificial neural networks. In this context, memory replay means that when a network is trained on new data, data of previous classes is added to the training data. It is customary to divide memory replay into two methods – rehearsal, in which training includes old examples of familiar classes, simulating seeing familiar people again, and generative replay, in which training includes recalled samples from memory, as may happen during visual imagery or sleep. Thus, combining rehearsal and generative replay effectively makes continual training similar to batch-training. This principle has been implemented by Shin et al. (2017), who used a Generative Adversarial Network (GAN) trained on object categories from the CIFAR-10 dataset, and showed that previously learned knowledge can be preserved by generating synthetic examples of earlier classes, during the learning of new ones (Shin et al., 2017).

Building on these insights, we propose a three-module architecture that integrates principles of CLS and memory replay within a unified computational framework for face learning. The model consists of an **Embedder**, which gradually learns to construct identity representations for faces; an **Autoencoder**, which reconstructs realistic face images from stored embeddings, enabling generative replay; and a **Memory module**, which stores and retrieves embeddings of previously encountered individuals (see Figure 1). Together, these components embody a functional division reminiscent of CLS: the embedder captures the slow learning, generalizing properties of the neocortex, while the memory module supports fast storage of idiosyncratic episodes akin to the hippocampus, with the autoencoder serving as the replay mechanism that links the two. This architecture allows continual learning of new identities while preserving previously acquired knowledge, thereby offering a biologically grounded account of human-like face learning over time.

**Figure 1.**
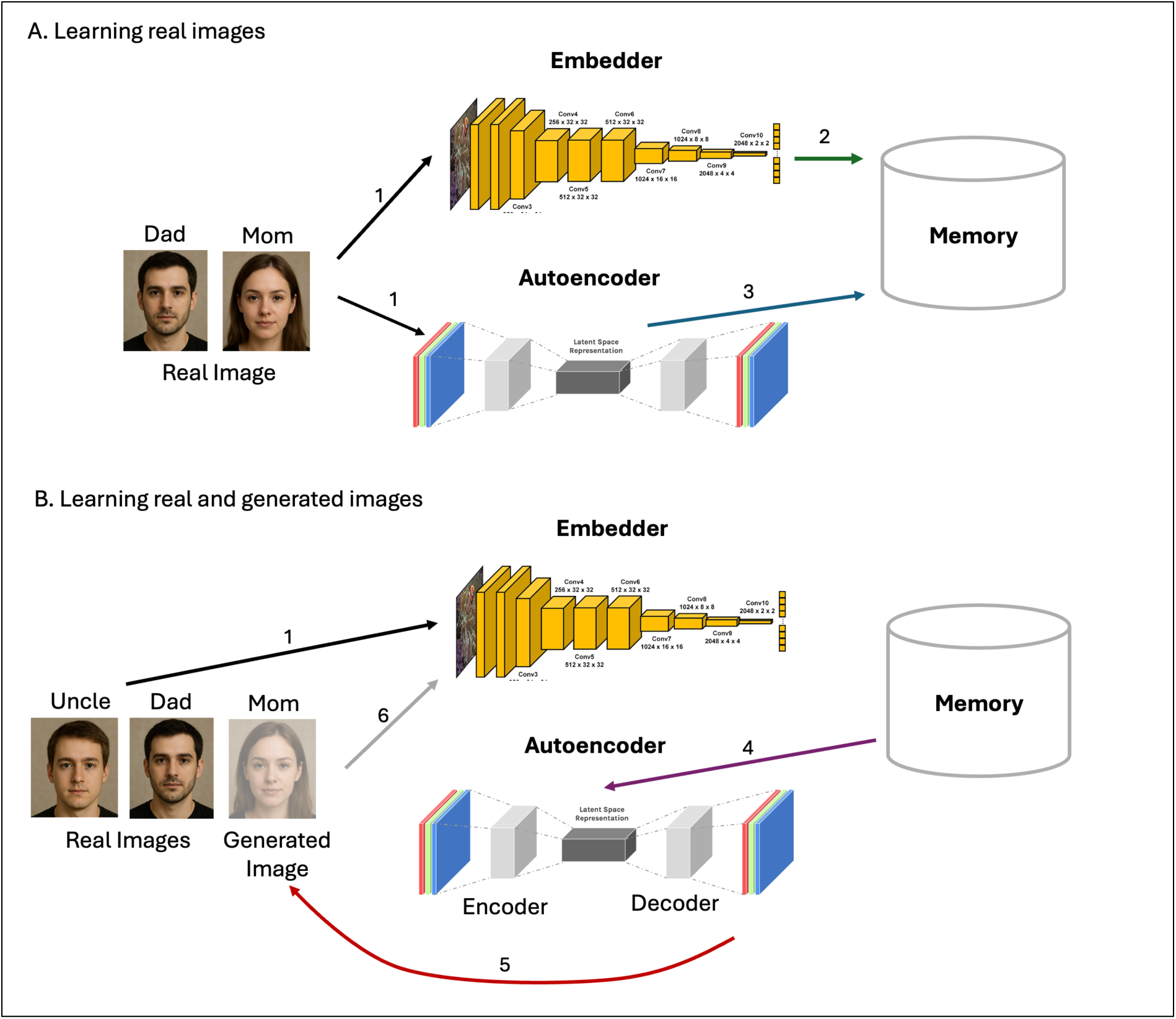
A: 1. The embedder and the autoencoder are trained on real face images (e.g., mom and dad) (black arrows). 2. The embeddings of the faces generated by the embedder are stored in memory for recognition (green arrow). 3. The embeddings of the faces generated by the autoencoder bottleneck are stored in memory for replay (blue arrow). B: 4. The autoencoder uses the embeddings of images of familiar identities stored in memory for replay (purple arrow) 5. Images of familiar faces are generated by the decoder (red arrow) 6. The generated images are used for training the embedder on identities that are absent (e.g., mom) (gray arrow) in addition to new (e.g., uncle) and old (e.g., dad) identities that are present (black arrows).

To support effective replay and recognition of familiar identities, our memory module is built using a multiple prototype architecture. Previous computational models of face recognition often store a single averaged representation for each identity (Burton et al., 2005). Although such averaging supports generalization across different images of the same person, it can lead to high false recognition rates (Noyes et al., 2021) and may produce representations that do not correspond to any real appearance. For instance, averaging images before and after a major change in hairstyle may yield an intermediate, non-existent representation. To address this, our approach draws inspiration from the **SUSTAIN** model of category learning (Love et al., 2004), which represents categories through multiple prototypes, formed by clustering exemplars based on their visual similarity. Similarly, our memory module represents each identity as a set of multiple clusters in the embedding space, each capturing a subset of within-identity perceptually similar face images. Cluster centroids serve as compact exemplars, while local variability around them preserves realistic appearance diversity. This multiple-prototype structure offers a biologically plausible balance between memory capacity and representational richness: it avoids storing every individual instance while retaining the flexibility needed to recognize familiar faces across substantial visual variation. A main advantage of this multiple prototype representation is that during generative replay, reconstructing these cluster centroids and sampling around them produces high-quality, variable synthetic faces required for successful continual learning.

Using this multiple-prototype memory structure, face recognition in our model is implemented through a **similarity-based mechanism**: the embedding of an incoming face is compared to the stored embeddings of familiar identities in memory, and recognition is assigned to the most similar cluster. This approach follows proposals that human face recognition operates via similarity-based retrieval (Kramer et al., 2018; Noyes et al., 2021).

Our model’s design, separating between the Embedder and Memory, introduces a functional distinction between representation learning and person recognition, closely paralleling the CLS framework: the Embedder plays the role of the neocortex, gradually constructing a generalizable representational space for faces, while the Memory module functions analogously to the hippocampus, storing specific exemplars without further modification of network weights. Such separation allows the model to acquire new faces instantaneously, simply by inserting their embeddings into memory, mirroring the human ability to recognize a person after a single encounter.

Within this framework, a face is considered *familiar* if its representation exists in memory, regardless of whether it was part of the embedder’s original training set. In addition, the SUSTAIN-inspired multiple-prototype memory structure may account for the differing robustness of familiar and unfamiliar face recognition (Jenkins et al., 2011; Kramer et al., 2017; Young & Burton, 2018). Familiar identities, represented by several clusters, can be matched across wide appearance variations such as aging or hairstyle changes, whereas unfamiliar faces rely on surface-level similarity that may be generalized well within prototypes but not across different prototypes of the same identity.

Finally, the separation between the embedder, which generates face representations, and memory, which stores familiar identities, allows a gradual stabilization of the embedder, which may model a developmental critical period (McKone et al., 2019; Pascalis et al., 2020): The importance of experience with faces during the first year of life was demonstrated in individuals who were treated for cataract removal during the first year of life, who showed face recognition impairments in adulthood (Gilad-Gutnick et al., 2024; Maurer, 2017). This can be implemented in our model where early experiences shape the representational geometry generated by the embedder, after which recognition depends primarily on memory retrieval. Nonetheless, continued exposure may still fine-tune representations, consistent with lifelong but reduced cortical plasticity.

To anchor our model within a realistic developmental framework, we sought to emulate the conditions under which an infant’s face recognition system emerges. This requires addressing not only the challenge of continual learning, but also the ecological constraints of early visual experience. Studies with head mounted cameras on infants’ foreheads documented the statistics of their face diet during the first year of life (Fausey et al., 2016; Jayaraman et al., 2015). These data show that infants encounter a very limited number of individuals during the first year of life. Furthermore, the visual input is confined to naturalistic, dynamic contexts. To approximate these conditions, we trained our model on a small set of identities (up to ten), using images cropped from a television series to simulate the continuous and coherent exposure characteristic of real-world viewing (see *Methods* for details). In the following section, we describe the architecture of the model and the procedures used for training and testing.

## Methods

### Model Architecture

Our face recognition model consists of three components: an Embedder, an Autoencoder, and a Memory module (see Figure 1). Together, these components support incremental acquisition and retention of face identities through continual learning with memory replay.

### Embedder

The Embedder is a deep convolutional neural network (DCNN) trained to generate face representations optimized for identity discrimination (Figure 1A.1). In this study, we adopt the ArcFace model proposed by (Deng et al., 2019), which is based on the ResNet architecture. The network produces direct embeddings and is trained using an angular margin-based loss function that enhances inter-class separability and intra-class compactness. The network is trained “from scratch”, starting with random weights. Network details and parameters are described in the Supplementary Materials.

### Autoencoder

The Autoencoder consists of an Encoder and a Decoder, forming a symmetric convolution-deconvolution architecture. The Encoder compresses the input image into a low-dimensional latent representation (“bottleneck”), while the Decoder reconstructs the image from this representation. The network is trained end-to-end to minimize the pixel-wise reconstruction error between input and output images, thereby learning to generate realistic reconstructions from compact codes. Like the Embedder, the Autoencoder is also trained “from scratch”. Network details and parameters are described in the Supplementary Materials.

## Memory

### Multiple-prototype memory construction via clustering

For each identity, we extract all embedding vectors from the corresponding training images. These embeddings are clustered using agglomerative clustering, a hierarchical, non-parametric method, that does not require a predefined number of clusters. This flexibility allows memory structure to naturally reflect the intrinsic variability of the face representations of each identity. For each resulting cluster, we compute and store two statistical descriptors:

#### Centroid

Rather than computing the mean of the cluster, we identify the centroid as the embedding vector closest to the cluster mean. This preserves the biological plausibility of storing an actual exemplar rather than an abstract average.

#### Standard Deviation (σ)

We compute the standard deviation of vectors within each cluster to characterize the local spread of representations.

Our framework includes two memory modules, corresponding to the two complementary forms of face representation: one derived from the Embedder (Figure 1A.2) and used for recognition, and the other derived from the Autoencoder (Figure 1A.3) and used for generative replay. This separation reflects a functional distinction between representational goals. The representations produced by the Embedder are optimized for identity discrimination—maximizing separability between individuals while minimizing within-identity variation. In contrast, the representations encoded by the Autoencoder are optimized for faithful image reconstruction, preserving fine-grained variability across dimerent views and conditions of the same identity. Attempting to satisfy both objectives within a single representational space leads to unstable training and suboptimal performance, as the two loss functions impose opposing constraints. This dual-route solution to this representational trade-om may point to a general computational principle and may be consistent with studies that suggest distinct neural mechanisms for perception and imagination (Breedlove et al., 2020; Lee et al., 2012). Both memory modules are constructed using the same procedure, and store representations in a multiple-prototype (cluster) structure. However, Autoencoder memory is expected to have more clusters per identity relative to the embedder, due to the dimerent nature of the training goals/objectives.

### Image Generation from Memory

The Autoencoder memory, constructed using bottleneck embeddings, supports generative replay of identities during continual learning (Figure 1A.4). To regenerate images for a given identity, the centroid embedding of each cluster is decoded into a face image (Figure 1A.5). Additional variability is introduced by sampling from a multivariate normal distribution centered on each centroid with variance given by σ_cluster_. The number of synthetic examples generated per cluster is proportional to its size during training, such that the distribution of replayed images reflects the original frequency of cluster membership. This process enables the replay of plausible, diverse examples of previously learned identities (Figure 1A.6).

### Recognition from Memory

Recognition of a new face is performed via a nearest-neighbor search. The test image is first passed through the Embedder to obtain its embedding, which is then compared to all cluster centroids using a normalized distance metric:

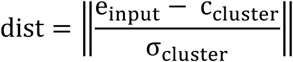

where e_input_ is the test embedding, c_cluster_ is a cluster centroid (both are normalized to unit length), and σ_cluster_is the cluster’s standard deviation. This normalization provides greater tolerance for clusters with high internal variability, while tightening the decision boundary around more homogeneous clusters. Recognition proceeds by selecting the identity associated with the nearest cluster. To categorize unfamiliar faces, a distance threshold is applied: if the minimum distance exceeds this threshold, the input is classified as “unfamiliar,” consistent with human performance in novelty detection.

### Matching unfamiliar faces

Although our model is focused on recognition of familiar faces, as this is the main goal of face recognition in infants, similar to human face system, it also supports matching of unfamiliar faces. To decide if two unfamiliar face images belong to the same identity or not, both images are run through the Embedder and the produced representations are compared for perceptual similarity (this procedure can be used also when one face is familiar and one is not).

### Model training

General Training Protocol:

1. Incremental identity exposure Training was organized into a series of learning steps, each introducing an increasing number of unique identities. Step 1 contained 2 identities, step 2 contained 3 identities, step 3 contained 4, and so on. This gradual exposure simulates the natural progression of person recognition in social settings, where new individuals are encountered over time.
2. Substep-based identity combinations Each step was further divided into substeps, with each substep comprising a subset of 2 identities drawn from the current step. For example, in step 2 (identities 1–3), substeps included the combinations [1,2], [1,3], and [2,3]. This structure reflects realistic social conditions in which learners rarely encounter all known identities simultaneously (e.g., in dyadic or triadic interactions). To control for order effects and ensure balanced exposure, the order of substeps within each step was randomized across training runs.
3. Joint training of Autoencoder and Embedder During each substep, both the Embedder and Autoencoder were trained simultaneously. The training process for each substep was initialization-aware: The Embedder for a given substep was initialized from the trained weights of the previous substep and trained for up to 100 epochs, or until the mean training accuracy reached 100%, whichever came first. The Autoencoder was similarly initialized from its state at the previous substep, but trained for a fixed 20 epochs per substep. At the conclusion of each substep, the Memory module was updated with representations generated by the current models.
4. Generative Replay to Prevent Catastrophic Forgetting Because each substep included only a subset of the current step’s identities, before training the Embedder in each substep, the stored bottleneck codes from previous substeps were decoded by the Autoencoder to regenerate images of missing identities. These synthetic reconstructions were then included alongside the real images in the training batch. For example, if a substep involved only identities [1,3], synthetic images of identity 2 were generated and included in training.

To train our model, we used two protocols:

<u>Protocol 1:</u> Incremental Exposure to 10 Identities The model was trained on a total of 10 unique identities, introduced sequentially across nine learning steps, starting from 2 identities and up to 10 identities. This protocol was designed to test the most difficult scenario in which the number of generated images is maximized relative to real ones.
<u>Protocol 2:</u> Naturalistic Exposure inspired by Infant Head-mounted Camera Data The second protocol was designed to mirror more closely empirical findings from infant experience, based on Smith and colleagues, who used head-mounted cameras to quantify the visual exposure of infants to faces during the first year of life (Fausey et al., 2016; Jayaraman et al., 2015). Their findings indicated that infants are typically exposed to a total of ∼8 individuals, and that most face-containing scenes include a small, stable set of ∼3 familiar identities. Accordingly, in this second protocol, learning steps began with real images of 3 identities and increased incrementally up to 8. Each step was again divided into randomized substeps, but substeps now consisted of combinations of real images of 3 or 4 identities, rather than pairwise groupings. This structure better approximates the stable social environment described in developmental studies. Thus, unlike Protocol 1, there was a subset of 3-4 identities that were always visible via real images in each substep; the remaining identities were recalled via generative memory replay using the Autoencoder and Memory module.

### Training Images

To reflect the natural within-identity variability that is accumulated from dynamic continuous visual input, that characterizes early visual experience, we constructed a dataset with constrained diversity. Specifically, we extracted face images from television series, where individual characters maintain a relatively stable appearance across episodes. For each identity, 1,000 images were collected by automatically cropping and aligning face regions using face detection and classification (DeepFace, https://github.com/serengil/deepface). The resulting dataset contained natural variation in pose, illumination, and facial expression, but minimal variation in age or hairstyle. This mirrors more closely the type of consistent visual exposure infants receive during interactions with familiar caregivers. The full dataset contained images of 20 characters. We repeated the model training and evaluation 10 times, each time randomly selecting 10 identities (in Protocol 1) or 8 identities (in Protocol 2), and randomly ordering their appearance. Model evaluation results (see Results section) are shown as mean (and standard deviation) of the 10 repetitions.

### Recognition Evaluation based on familiar identities

As mentioned above, our model is designed to simulate the goal of human face recognition, which is to identify familiar individuals within social contexts. Accordingly, our evaluation focused on the model’s ability to recognize previously encountered identities. Recognition performance was assessed after each learning step (e.g., following the acquisition of 2, 3, 4, or more identities). At each of these checkpoints, we evaluated the model’s recognition for all identities learned up to that point. To this end, we selected test images of these same identities that had not been included in training. These novel images were processed by the Embedder to generate test embeddings, which were matched against the stored identity clusters in memory using a nearest-neighbor approach. The identity of the closest cluster (based on normalized distance) was taken as the recognition output. Accuracy was defined as the proportion of test images for which the predicted identity matched the ground truth, across all known identities at each step.

### Unfamiliar face matching evaluation

In addition to testing our model on familiar faces, we tested the model’s performance on unfamiliar faces, using the Labeled Faces in the Wild (LFW) benchmark (Huang et al., 2008). This also allowed us to compare our model to existing face recognition models (e.g., Taigman et al., 2014).

### Comparison training modes

To contextualize the performance of our continual learning with replay model (“CL w/replay mode”), we evaluated it alongside two alternative training modes: a batch training mode (“Batch mode”) and a naive continual learning mode, without memory replay (“Naive CL mode”). These comparisons were designed to isolate the contributions of memory-based replay and incremental learning structure.

#### Batch mode

The Batch mode served as an upper-bound benchmark for recognition performance. Like the CL w/replay mode, it progressed through learning steps containing 2, 3, 4, and more identities. However, instead of incremental exposure, in Batch mode the Embedder was trained on all identities in each step simultaneously. Similar to the CL w/replay mode, the Embedder at each step was initialized from the trained model of the previous step, allowing knowledge accumulation over time.

#### Naive Continual learning mode, without memory replay

The second comparison was designed to isolate the effect of memory replay in preventing catastrophic forgetting. This mode used the same stepwise and substep structure as the CL w/replay mode, with incremental exposure to identities, but no generative replay was used—each substep only included images from the currently visible identities, and no reconstructions of previously seen identities were included. As a result, the model was susceptible to interference and forgetting. As in the other modes, the Embedder at each step was initialized from the previous step’s model to simulate cumulative learning. This mode is expected to exhibit catastrophic forgetting, manifested as declining recognition accuracy for previously learned identities across training steps.

We used the same recognition evaluation procedure described above also for the Batch and the Naïve CL models.

## Results

### Quality of the reconstructed face images

We first examined the image quality of the faces generated by the Autoencoder of the trained identities, across the different stages of learning. Figure 2 shows the mean reconstruction error, calculated as follows: we ran the trained images of the identities of each step through the autoencoder and measured the mean squared error (MSE) between input and reconstructed images. This is not a direct measure of the quality of the generated images that are used during training, since those images were generated from stored embeddings of prototypes (the cluster centroids) and from random samples around them. However, this is a measure of the improvement of the autoencoder, which directly determines the quality of the generated images. Results show that reconstruction error decreases reaching a minimal level when the autoencoder is trained on 6 identities or more.

**Figure 2.**
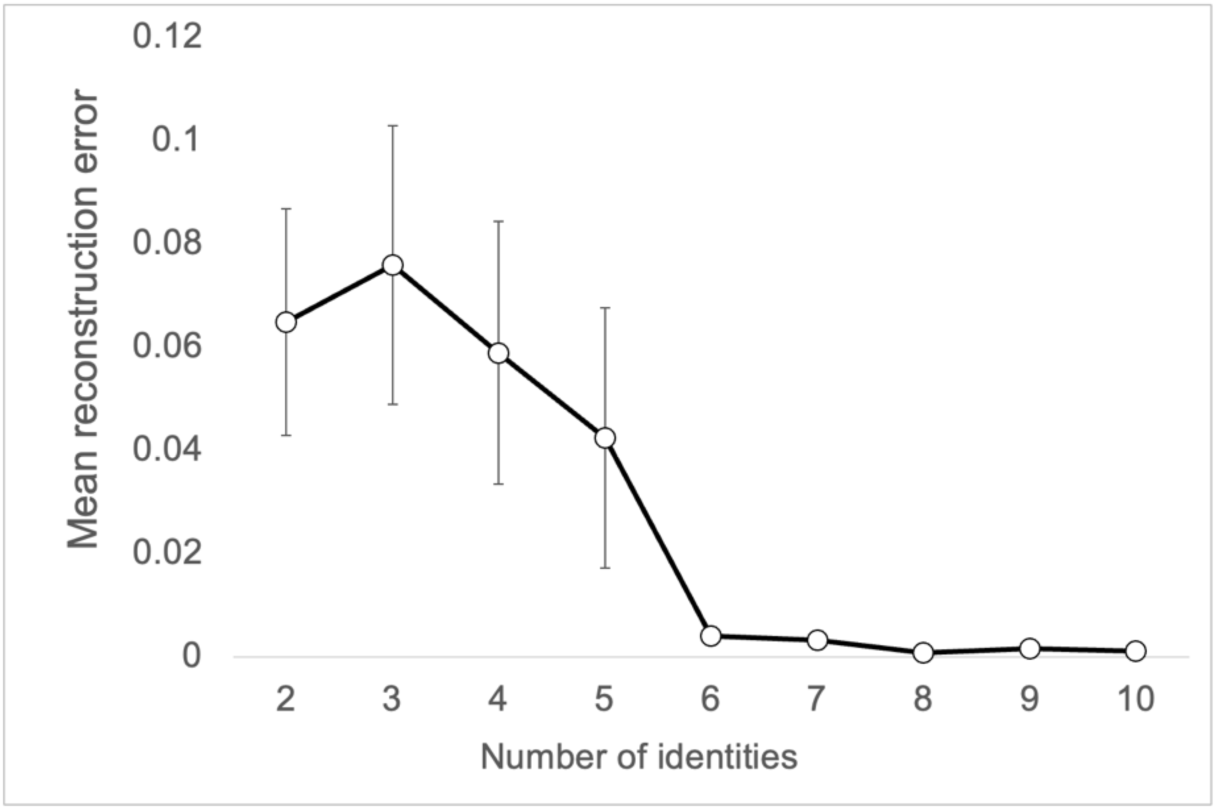
The mean and standard deviation reconstruction error of the reconstructed images at each step of training of the autoencoder, indicating that the quality of the face images improved as the autoencoder was trained on more identities.

### Recognition Performance for trained (familiar) faces

The two training protocols revealed very similar findings. Figure 3 shows mean and standard error of recognition rates (y-axis) for the 3 training modes, across 10 samples of trained identities. The x-axis indicates the number of trained identities at each step, starting from training on 2 identities up to 10 identities in protocol 1 (left) and starting from 3 identities up to 8 identities in protocol 2 (right). Performance for the Batch mode showed the best performance across all training steps. Performance for the Naïve CL mode decreased as additional identities were added to the training set, reflecting catastrophic forgetting. Performance of the CL w/replay mode was initially low like the Naïve CL mode, but then improved to the level of the Batch mode when more than 5 identities were included in the training set. Performance for the Naïve CL mode was overall better in protocol 2 than protocol 1 because three identities were repeatedly presented from their real images.

**Figure 3:**
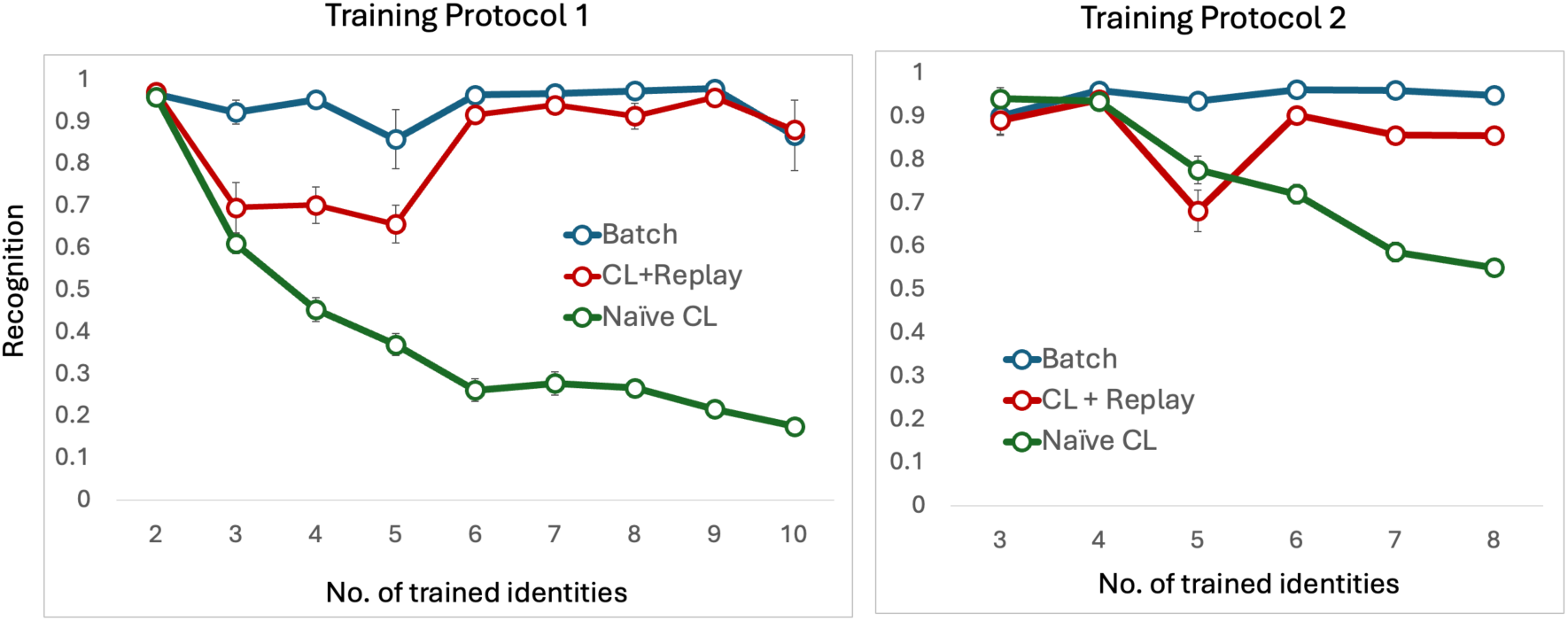
Recognition (proportion correct) performance of untrained images of familiar identities for the three training modes in training protocols 1 & 2 as a function of the number of trained identities.

*Training protocol 1*: A repeated measure ANOVA with number of trained IDs (2-10 identities) and training mode (Batch, Naïve CL, CL w/replay) revealed a main effect of number of IDs F(8,216) = 29.6, p < .001, partial η^+^= 0.52, a main effect of training mode F(2,27) = 561.67, p < .001, partial η^+^= 0.98 and an interaction between number of IDs and training mode, F(8,216) = 27.31, p < .001, partial η^+^= 0.67). Post hoc comparisons within each training step (see supplementary Table 1) show that the CL w/replay mode was initially significantly worse than the Batch mode but not from the Naïve CL mode. When training increased to 6 identities, performance of CL w/replay mode was better than the Naïve mode and did not differ statistically from the Batch mode. This improved performance is in line with the decrease in reconstruction error of the generated images (see Figure 2).

*Training protocol 2*: A repeated measure ANOVA with number of trained IDs (3-8 identities) and training mode (Batch, Naïve CL, CL w/replay) as two factors. There was a main effect of number of IDs F(5,125) = 23.78, p < .001, partial η^+^= 0.47, a main effect of training mode F(2,27) = 84.3, p < .001, partial η^+^= 0.86 and an interaction between number of IDs and training mode, F(10,135) = 20.84, p < .001, partial η^+^= 0.6). Post hoc comparisons within each training step (see supplementary Table 2) indicate that performance did not differ between the three training modes during the initial stage of training when only 3 or 4 identities were learned. When learning increased to 5 identities, performance on the CL w/replay and Naïve CL modes was significantly lower than the Batch mode and did not differ from one another. Once 6 identities or more were learned, performance on the CL w/replay mode was higher than the Naive CL mode and did not differ statistically from the Batch mode.

### Verification performance for untrained identities

In addition to testing recognition of familiar faces, we also tested our model for matching unfamiliar faces, using the LFW benchmark. This also enabled us to compare our model to existing face recognition DCNNs, which are typically evaluated based on performance on untrained identities. Figure 4 shows that as the number of train identities increased, performance improved for Batch and CL w/replay modes, but not for the Naive CL mode.

**Figure 3:**
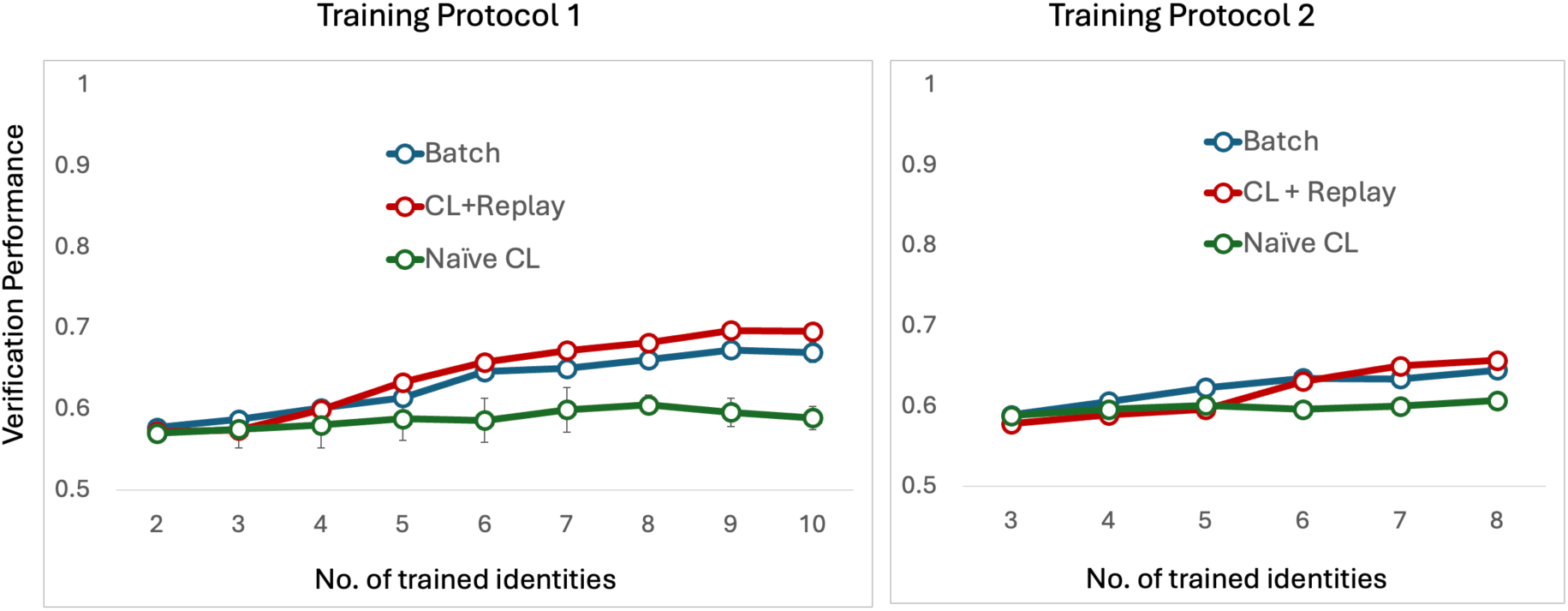
Verification performance for the three training modes for identity matching of untrained identities in training protocols 1 & 2, as a function of the number of trained identities.

*Training protocol 1*: A repeated measure ANOVA with number of trained IDs (2-10 identities) and training mode (Batch, Naïve CL, CL w/replay) revealed a main effect of number of IDs F(8,216) = 80.41, p < .001, partial η^+^= 0.75, a main effect of training mode F(2,27) = 32.54, p < .001, partial η^+^= 0.71 and an interaction between number of IDs and learning mode, F(16,216) = 11.32, p < .001, partial η^+^= 0.46). Post hoc comparisons (see supplementary Table 3) show that starting from a training set of 6 identities there was no significant difference between the CL w/replay and Batch modes that were both significantly better than the Naïve CL mode.

*Training protocol 2*: An ANOVA with number of trained IDs (3-8 identities) and learning mode (Batch, Naïve CL, CL w/replay) as two factors. There was a main effect of number of IDs F(5,135) = 51.14, p < .001, partial η^+^= 0.65, a main effect of learning mode F(2,27) = 5.74, p = .008, partial η^+^= 0.30 and an interaction between number of IDs and learning mode, F(10,135) = 10.13, p < .001, partial η^+^= 0.43). Post hoc comparisons within each training step (see Supplementary Table 4) show that when training increased to 6 identities or more performance for CL w/replay and Batch modes did not differ and both were better than the Naïve CL mode.

## Discussion

Current computational models of face recognition achieve human-level accuracy and reproduce several human-like face effects (Dobs et al., 2023; Phillips & White, 2025; Yovel et al., 2023), yet they learn in ways that differ fundamentally from human development. We propose a framework that bridges this gap by aligning the learning dynamics of artificial networks with the ecological and developmental conditions under which humans acquire face recognition. Our model learns continually rather than in batches, is exposed to a small, socially realistic set of identities—consistent with infants’ early experience of only a few familiar caregivers (Kelly et al., 2007; Smith & Slone, 2017) and receives visual input that simulates temporally continuous, low-variability input. Centered on the premise that human face recognition primarily serves to identify familiar individuals, the model was evaluated on new images of learned faces and compared to two baselines: a batch-trained model and a naïve continual learning model without memory replay. As expected, the batch model achieved ceiling performance, while the naïve continual learner suffered from catastrophic forgetting (see Figure 3). Our model initially showed a modest decline, due to the low quality of early replay images (see Figure 2) but quickly recovered as the quality of the generated images improved, reaching the batch model’s high accuracy after learning 4–5 identities across two training protocols.

We also evaluated our model on unfamiliar face verification, using the Labeled Faces in the Wild (LFW) benchmark. The continual learning with replay model and the batch model achieved a moderate level of performance of about 70% accuracy, despite being trained on only a very small dataset (up to 10 identities × 1,000 images each). While this accuracy is lower than state-of-the-art deep convolutional networks, it reflects realistic developmental constraints and may approximate human performance in unfamiliar face recognition (Ritchie et al., 2015; Yovel et al., 2024).

Our framework provides a perspective on the distinction between familiar and unfamiliar face processing. We begin from the premise that the primary function of the human face recognition system, particularly during development, is to identify familiar individuals within a social context, rather than to match the identities of unfamiliar faces. Notably, image-matching tasks themselves are not ecologically natural, as they became possible only with the advent of photographic media. Thus, when modelling the developmental trajectory of face recognition, it is plausible to assume that early learning mechanisms evolved to support the recognition of socially relevant, familiar individuals rather than image-based face comparison. Moreover, behavioral research has consistently shown that humans are markedly better at matching images of familiar faces than unfamiliar ones, especially under conditions of appearance variation such as changes in lighting, pose, or age (Jenkins et al., 2011; Burton, 2013; Ritchie & Burton, 2017)

Several computational accounts have attempted to model this discrepancy between performance level for familiar and unfamiliar faces (Blauch et al., 2021; Kramer et al., 2018; Noyes et al., 2021). Our model differs from these models in the following ways (see supplementary Table 5 for a detailed comparison between the models): First, as aforementioned, current models are trained in a batch mode, while our model is trained continually. Second, current models assume different perceptual representation of familiar and unfamiliar faces. For example, Kramer et al (2018) suggested that unfamiliar faces are represented by a PCA while familiar faces by a PCA + LDA. In Blauch et al (2021), unfamiliar faces are represented by the penultimate layer and familiar faces by the output layer of a DCNN. In our model, which does not involve retraining for familiar faces, the perceptual representation of familiar and unfamiliar faces is similar. The advantage in recognition of familiar over unfamiliar faces is due to their representation in the memory module that enables identification of the familiar identities, which is not available for unfamiliar faces. Thus, our model separates representation learning and memory storage: the embedder provides a stable feature space analogous to cortical representations, while the memory module stores representations of familiar identities directly. This division enables memory-based acquisition of new faces without retraining, providing a more biologically plausible account of how familiar face representations may accumulate over the lifespan. Finally, while previous models represented familiar faces in memory as averages (Burton et al., 2005; Noyes et al., 2021), familiar faces in our model are represented in memory by multiple prototypes, rather than a single prototype for each identity. This supports both generalization and discrimination for highly variable images of same and different identities.

While our architecture draws inspiration from the Complementary Learning Systems (CLS) framework, its contribution lies in extending these principles into the domain of face learning and recognition, providing a concrete demonstration of how CLS-like mechanisms can support human-like recognition behavior, in a perceptual domain that requires both flexibility and precision. In classical CLS theory, the hippocampus and neocortex operate as two learning systems with distinct but interacting roles: the hippocampus rapidly encodes unique experiences, whereas the neocortex gradually extracts structured regularities through slow integration across episodes (Kumaran et al., 2016; McClelland et al., 1995). Our results illustrate how this division of labor can be computationally instantiated: The embedder–memory distinction captures the trade-om between generalization and individuation – the embedder (analogous to the neocortex) organizes perceptual regularities across faces, while the memory module (analogous to the hippocampus) retains the distinctive signatures of known individuals. Generative replay, implemented through the autoencoder, operationalizes the CLS idea of hippocampal reactivation as a training signal for cortical consolidation. Overall, this model omers a substantial new computational framework to face learning and recognition, in contrast to existing face recognition models that include only a face-trained embedder and are therefore limited to batch learning of all known faces at once.

Our model can also implement effects of *critical periods* in face perception (Gilad-Gutnick et al., 2024; Maurer, 2017). The embedder’s early stabilization parallels cortical specialization for faces during infancy, while the continued adaptability of the memory module may model how adults can still acquire new familiar faces without restructuring their representational space. Thus, our framework situates face learning within a continuous developmental trajectory—from early structural tuning to mature episodic flexibility and can be used in future studies to study adult level face recognition as well as other robust face recognition phenomenon such as the other race effect (McKone et al., 2019).

In summary, by integrating continual learning, ecologically realistic training conditions, and a separation between representational learning and identity memory, our model propose a new computational framework that provides a more biologically grounded account of how humans acquire, store, and recognize faces across the lifespan.

## References

1. Abudarham, N., Shkiller, L., & Yovel, G. (2019). Critical features for face recognition. Cognition, 182, 73–83. 10.1016/j.cognition.2018.09.002

2. Blauch, N. M., Behrmann, M., & Plaut, D. C. (2021). Computational insights into human perceptual expertise for familiar and unfamiliar face recognition. Cognition, 208, 104341. 10.1016/j.cognition.2020.104341

3. Breedlove, J. L., St-Yves, G., Olman, C. A., & Naselaris, T. (2020). Generative Feedback Explains Distinct Brain Activity Codes for Seen and Mental Images. Current Biology, 30(12), 2211–2224.e6. 10.1016/j.cub.2020.04.014

4. Burton, A. M., Jenkins, R., Hancock, P. J. B., & White, D. (2005). Robust representations for face recognition: The power of averages. Cognitive Psychology, 51(3), 256–284. 10.1016/j.cogpsych.2005.06.003

5. Carr, M. F., Jadhav, S. P., & Frank, L. M. (2011). Hippocampal replay in the awake state: A potential substrate for memory consolidation and retrieval. Nature Neuroscience, 14(2), 147–153. 10.1038/nn.2732

6. Deng, J., Guo, J., Xue, N., & Zafeiriou, S. (2019). ArcFace: Additive Angular Margin Loss for Deep Face Recognition. 4690–4699. https://openaccess.thecvf.com/content_CVPR_2019/html/Deng_ArcFace_Additive_Angular_Margin_Loss_for_Deep_Face_Recognition_CVPR_2019_paper.html

7. Dobs, K., Yuan, J., Martinez, J., & Kanwisher, N. (2023). Behavioral signatures of face perception emerge in deep neural networks optimized for face recognition. Proceedings of the National Academy of Sciences, 120(32), e2220642120. 10.1073/pnas.2220642120

8. Fausey, C. M., Jayaraman, S., & Smith, L. B. (2016). From faces to hands: Changing visual input in the first two years. Cognition, 152, 101–107. 10.1016/j.cognition.2016.03.005

9. French, R. M. (1999). Catastrophic forgetting in connectionist networks. Trends in Cognitive Sciences, 3(4), 128–135. 10.1016/S1364-6613(99)01294-2

10. Gilad-Gutnick, S., Hu, H. F., Dalrymple, K. A., Gupta, P., Shah, P., Ralekar, C., Verma, D., Tiwari, K., Ben-Ami, S., Swami, P., Ganesh, S., Mathur, U., & Sinha, P. (2024). Face-specific identification impairments following sight-providing treatment may be alleviated by an initial period of low visual acuity. Scientific Reports, 14(1), 17374. 10.1038/s41598-024-67949-z

11. Grossman, S., Gaziv, G., Yeagle, E. M., Harel, M., Mégevand, P., Groppe, D. M., Khuvis, S., Herrero, J. L., Irani, M., Mehta, A. D., & Malach, R. (2019). Convergent evolution of face spaces across human face-selective neuronal groups and deep convolutional networks. Nature Communications, 10(1), 4934. 10.1038/s41467-019-12623-6

12. Huang, G. B., Mattar, M., Berg, T., & Learned-Miller, E. (2008, October). Labeled Faces in the Wild: A Database forStudying Face Recognition in Unconstrained Environments. Workshop on Faces in “Real-Life” Images: Detection, Alignment, and Recognition. https://inria.hal.science/inria-00321923

13. Jayaraman, S., Fausey, C. M., & Smith, L. B. (2015). The Faces in Infant-Perspective Scenes Change over the First Year of Life. PLOS ONE, 10(5), e0123780. 10.1371/journal.pone.0123780

14. Jenkins, R., White, D., Van Montfort, X., & Burton, A. M. (2011). Variability in photos of the same face. Cognition, 121(3), 313–323.

15. Kelly, D. J., Liu, S., Ge, L., Quinn, P. C., Slater, A. M., Lee, K., Liu, Q., & Pascalis, O. (2007). Cross-Race Preferences for Same-Race Faces Extend Beyond the African Versus Caucasian Contrast in 3-Month-Old Infants. Infancy, 11(1), 87–95. 10.1080/15250000709336871

16. Khaligh-Razavi, S.-M., & Kriegeskorte, N. (2014). Deep Supervised, but Not Unsupervised, Models May Explain IT Cortical Representation. PLoS Computational Biology, 10(11), e1003915. 10.1371/journal.pcbi.1003915

17. Kramer, R. S., Jenkins, R., Young, A. W., & Burton, A. M. (2017). Natural variability is essential to learning new faces. Visual Cognition, 25(4–6), 470–476.

18. Kramer, R. S. S., Young, A. W., & Burton, A. M. (2018). Understanding face familiarity. Cognition, 172, 46–58. 10.1016/j.cognition.2017.12.005

19. Kumaran, D., Hassabis, D., & McClelland, J. L. (2016). What Learning Systems do Intelligent Agents Need? Complementary Learning Systems Theory Updated. Trends in Cognitive Sciences, 20(7), 512–534. 10.1016/j.tics.2016.05.004

20. Lee, S.-H., Kravitz, D. J., & Baker, C. I. (2012). Disentangling visual imagery and perception of real-world objects. NeuroImage, 59(4), 4064–4073. 10.1016/j.neuroimage.2011.10.055

21. Love, B. C., Medin, D. L., & Gureckis, T. M. (2004). SUSTAIN: A Network Model of Category Learning. Psychological Review, 111(2), 309–332. 10.1037/0033-295X.111.2.309

22. Maurer, D. (2017). Critical periods re-examined: Evidence from children treated for dense cataracts. Cognitive Development, 42, 27–36. 10.1016/j.cogdev.2017.02.006

23. McClelland, J. L., McNaughton, B. L., & O’Reilly, R. C. (1995). Why there are complementary learning systems in the hippocampus and neocortex: Insights from the successes and failures of connectionist models of learning and memory. Psychological Review, 102(3), 419–457. 10.1037/0033-295X.102.3.419

24. McCloskey, & Cohen. (1989). Catastrophic interference in connectionist networks: The sequential learning problem. In Psychology of learning and motivation. Academic Press., *24*, 109–165.

25. McKone, E., Wan, L., Pidcock, M., Crookes, K., Reynolds, K., Dawel, A., Kidd, E., & Fiorentini, C. (2019). A critical period for faces: Other-race face recognition is improved by childhood but not adult social contact. Scientific Reports, 9(1), 12820. 10.1038/s41598-019-49202-0

26. Mike Burton, A. (2013). Why has research in face recognition progressed so slowly? The importance of variability. Quarterly Journal of Experimental Psychology, 66(8), 1467–1485. 10.1080/17470218.2013.800125

27. Noyes, E., Parde, C. J., Colón, Y. I., Hill, M. Q., Castillo, C. D., Jenkins, R., & O’Toole, A. J. (2021). Seeing through disguise: Getting to know you with a deep convolutional neural network. Cognition, 211, 104611.

28. Ólafsdóttir, H. F., Bush, D., & Barry, C. (2018). The Role of Hippocampal Replay in Memory and Planning. Current Biology, 28(1), R37–R50. 10.1016/j.cub.2017.10.073

29. O’Toole, A. J., & Castillo, C. D. (2021). Face recognition by humans and machines: Three fundamental advances from deep learning. Annual Review of Vision Science, 7, 543–570.

30. Pascalis, O., Fort, M., & Quinn, P. C. (2020). Development of face processing: Are there critical or sensitive periods? Current Opinion in Behavioral Sciences, 36, 7–12. 10.1016/j.cobeha.2020.05.005

31. Phillips, P. J., & White, D. (2025). The state of modelling face processing in humans with deep learning. British Journal of Psychology, bjop.12794. 10.1111/bjop.12794

32. Ritchie, K. L., & Burton, A. M. (2017). Learning faces from variability. Quarterly Journal of Experimental Psychology, 70(5), 897–905.

33. Ritchie, K. L., Smith, F. G., Jenkins, R., Bindemann, M., White, D., & Burton, A. M. (2015). Viewers base estimates of face matching accuracy on their own familiarity: Explaining the photo-ID paradox. Cognition, 141, 161–169. 10.1016/j.cognition.2015.05.002

34. Schapiro, A. C., McDevitt, E. A., Rogers, T. T., Mednick, S. C., & Norman, K. A. (2018). Human hippocampal replay during rest prioritizes weakly learned information and predicts memory performance. Nature Communications, 9(1), 3920. 10.1038/s41467-018-06213-1

35. Shin, H., Lee, J. K., Kim, J., & Kim, J. (2017). Continual learning with deep generative replay. Advances in Neural Information Processing Systems, 30. https://proceedings.neurips.cc/paper/2017/hash/0efbe98067c6c73dba1250d2beaa81f9-Abstract.html

36. Smith, L. B., & Slone, L. K. (2017). A Developmental Approach to Machine Learning? Frontiers in Psychology, 8. 10.3389/fpsyg.2017.02124

37. Taigman, Y., Yang, M., Ranzato, M., & Wolf, L. (2014). DeepFace: Closing the Gap to Human-Level Performance in Face Verification. 1701–1708. https://openaccess.thecvf.com/content_cvpr_2014/html/Taigman_DeepFace_Closing_the_2014_CVPR_paper.html

38. van Dyck, L. E., & Gruber, W. R. (2023). Modeling Biological Face Recognition with Deep Convolutional Neural Networks. Journal of Cognitive Neuroscience, 35(10), 1521–1537. 10.1162/jocn_a_02040

39. Yamins, D. L. K., Hong, H., Cadieu, C. F., Solomon, E. A., Seibert, D., & DiCarlo, J. J. (2014). Performance-optimized hierarchical models predict neural responses in higher visual cortex. Proceedings of the National Academy of Sciences, 111(23), 8619–8624. 10.1073/pnas.1403112111

40. Yin, R. K. (1969). Looking at upside-down faces. Journal of Experimental Psychology, 81(1), 141–145. 10.1037/h0027474

41. Young, A. W., & Burton, A. M. (2018). Are We Face Experts? Trends in Cognitive Sciences, 22(2), 100–110. 10.1016/j.tics.2017.11.007

42. Yovel, G., Bash, E., & Bate, S. (2024). Humans’ extreme face recognition abilities challenge the well-established familiarity effect. Cognition, 251, 105904. 10.1016/j.cognition.2024.105904

43. Yovel, G., Grosbard, I., & Abudarham, N. (2023). Deep learning models challenge the prevailing assumption that face-like effects for objects of expertise support domain-general mechanisms. Proceedings of the Royal Society B: Biological Sciences, 290(1998), 20230093. 10.1098/rspb.2023.0093

