## Supplemental Material for "A developmentally inspired computational model of face recognition that learns continuously through generative memory replay"

Supplementary material

Methods

Supplementary Materials

### 1. Embedder implementation details:

For the Embedder model we based our code on the "arcface" project (https://github.com/ronghuaiyang/arcface-pytorch). The configuration parameters were as follows: resnet-18 backbone, num_classes 2000, arc_margin metric, focal_loss, sgd optimizer with an initial learning rate of 1e-1, learning_rate_decay of 0.95 every 10 steps, and weight_decay of 5e-4. In each training substep we started from the previous model, and trained up to 100 epochs, or until train accuracy reached 100% (which typically happened after ~20 epochs).

### 2. Autoencoder implementation details:

The Autoencoder was built from an Encoder and a Decoder. The Encoder included 5 [Conv2d-batchnormalization-leakyrelu-Dropout] layers, starting from image size of 128 by 128, and ended with a fully connected layer, outputting a bottleneck of size 100. The Decoder decoded the bottleneck via one fully connected layer followed by 5 [ConvTranspose2d-batchnormalization-leakyrelu] layers, reconstructing a 128 by 128 image. The Autoencoder was trained for 20 epochs in each training substep. Additional parameters were: MSE loss, learning rate - 0.0005, dropout – 0.4.

### 3. Memory construction details:

Both the recognition memory (storing Embedder representations) and the generative replay memory (storing Autoencoder bottlenecks) were constructed by clustering the representations of each identity. Clustering was performed using the *AgglomerativeClustering* algorithm from *scikit-learn* (metric = cosine, linkage = average). All representations were L2-normalized prior to clustering, and the distance threshold was set to 0.75. Owing to the different characteristics of the two representational spaces, the number and structure of clusters varied substantially between them. Embedder-derived embeddings were relatively homogeneous within each identity, yielding approximately 50 clusters per identity on average. In contrast, Autoencoder bottlenecks were more heterogeneous, producing around 200 clusters per identity. Cluster sizes were highly uneven in both memories: in the Embedder memory, a single large cluster typically contained ~750 representations, while most others were very small (median cluster size = 1.8). In the Autoencoder memory, the largest cluster contained ~60 representations (median = 2.9). Thus, although the overall clustering patterns differed between the two memories, both exhibited a similar structure in which some representations were isolated from all others. The resulting multi-prototype organization preserves this representational diversity, which might otherwise be lost in an averaged or single-prototype memory scheme.

### Results

Supplementary Table 1: Post hoc comparisons of the three training modes in each step in protocol 1 on recognition of familiar (trained) identities. CL: CL + Replay, Batch, Naïve: CL Naive

| no. trained IDs | Training mode | Training mode | Mean Difference | SE | t | Cohen's d | $p_{tukey}$ |
|---|---|---|---|---|---|---|---|
| 2 | CL | Batch | 0.005 | 0.045 | 0.112 | 0.05 | 1 |
| 2 | CL | Naive | 0.011 | 0.045 | 0.247 | 0.11 | 1 |
| 2 | Batch | Naive | 0.006 | 0.045 | 0.135 | 0.06 | 1 |
| 3 | CL | Batch | -0.227 | 0.045 | -5.091 | -2.277 | < .001 |
| 3 | CL | Naive | 0.086 | 0.045 | 1.929 | 0.863 | 1 |
| 3 | Batch | Naive | 0.313 | 0.045 | 7.02 | 3.14 | < .001 |
| 4 | CL | Batch | -0.251 | 0.045 | -5.63 | -2.518 | < .001 |
| 4 | CL | Naive | 0.248 | 0.045 | 5.562 | 2.488 | < .001 |
| 4 | Batch | Naive | 0.499 | 0.045 | 11.192 | 5.005 | < .001 |
| 5 | CL | Batch | -0.202 | 0.045 | -4.531 | -2.026 | 0.001 |
| 5 | CL | Naive | 0.287 | 0.045 | 6.437 | 2.879 | < .001 |

| | | | | | | | |
|---|---|---|---|---|---|---|---|
| 5 | Batch | Naive | 0.489 | 0.045 | 10.968 | 4.905 | < .001 |
| 6 | CL | Batch | -0.047 | 0.045 | -1.054 | -0.471 | 1 |
| 6 | CL | Naive | 0.655 | 0.045 | 14.691 | 6.57 | < .001 |
| 6 | Batch | Naive | 0.702 | 0.045 | 15.745 | 7.041 | < .001 |
| 7 | CL | Batch | -0.028 | 0.045 | -0.628 | -0.281 | 1 |
| 7 | CL | Naive | 0.662 | 0.045 | 14.848 | 6.64 | < .001 |
| 7 | Batch | Naive | 0.69 | 0.045 | 15.476 | 6.921 | < .001 |
| 8 | CL | Batch | -0.06 | 0.045 | -1.346 | -0.602 | 1 |
| 8 | CL | Naive | 0.647 | 0.045 | 14.511 | 6.49 | < .001 |
| 8 | Batch | Naive | 0.707 | 0.045 | 15.857 | 7.092 | < .001 |
| 9 | CL | Batch | -0.021 | 0.045 | -0.471 | -0.211 | 1 |
| 9 | CL | Naive | 0.741 | 0.045 | 16.62 | 7.433 | < .001 |
| 9 | Batch | Naive | 0.762 | 0.045 | 17.091 | 7.643 | < .001 |
| 10 | CL | Batch | 0.013 | 0.045 | 0.292 | 0.13 | 1 |
| 10 | CL | Naive | 0.705 | 0.045 | 15.812 | 7.071 | < .001 |
| 10 | Batch | Naive | 0.692 | 0.045 | 15.521 | 6.941 | < .001 |

Supplementary Table 2: Post hoc comparisons of the three training modes in each step in protocol 2 on recognition of familiar (trained) identities. CL: CL + Replay, Batch, Naïve: CL Naive

| # trained IDs | Training mode | Training mode | Mean Difference | SE | t | Cohen's d | $p_{tukey}$ |
|---|---|---|---|---|---|---|---|
| 3 | CL | Batch | -0.01 | 0.034 | -0.295 | -0.132 | 1 |
| 3 | CL | Naive | -0.05 | 0.034 | -1.477 | -0.66 | 0.991 |
| 3 | Batch | Naive | -0.04 | 0.034 | -1.181 | -0.528 | 0.999 |
| 4 | CL | Batch | -0.021 | 0.034 | -0.62 | -0.277 | 1 |
| 4 | CL | Naive | 0.004 | 0.034 | 0.118 | 0.053 | 1 |
| 4 | Batch | Naive | 0.025 | 0.034 | 0.738 | 0.33 | 1 |
| 5 | CL | Batch | -0.254 | 0.034 | -7.501 | -3.354 | < .001 |
| 5 | CL | Naive | -0.095 | 0.034 | -2.805 | -1.255 | 0.32 |
| 5 | Batch | Naive | 0.159 | 0.034 | 4.695 | 2.1 | < .001 |
| 6 | CL | Batch | -0.059 | 0.034 | -1.742 | -0.779 | 0.955 |
| 6 | CL | Naive | 0.182 | 0.034 | 5.375 | 2.404 | < .001 |
| 6 | Batch | Naive | 0.241 | 0.034 | 7.117 | 3.183 | < .001 |
| 7 | CL | Batch | -0.104 | 0.034 | -3.071 | -1.373 | 0.181 |
| 7 | CL | Naive | 0.27 | 0.034 | 7.973 | 3.566 | < .001 |
| 7 | Batch | Naive | 0.374 | 0.034 | 11.045 | 4.939 | < .001 |
| 8 | CL | Batch | -0.093 | 0.034 | -2.746 | -1.228 | 0.358 |
| 8 | CL | Naive | 0.305 | 0.034 | 9.007 | 4.028 | < .001 |

| | | | | | | | |
|---|---|---|---|---|---|---|---|
| 8 | Batch | Naive | 0.398 | 0.034 | 11.753 | 5.256 | < .001 |

Supplementary Table 3: Post hoc comparisons of the three training modes in each step in protocol 1 on the LFW benchmark. CL: CL + Replay, Batch, Naïve: CL Naive

| # trained IDs | Training mode | Training mode | Mean Difference | SE | t | Cohen's d | $p_{tukey}$ |
|---|---|---|---|---|---|---|---|
| 2 | CL | Batch | -0.005 | 0.011 | -0.455 | -0.204 | 1 |
| 2 | CL | Naive | 0.002 | 0.011 | 0.182 | 0.081 | 1 |
| 2 | Batch | Naive | 0.007 | 0.011 | 0.637 | 0.285 | 1 |
| 3 | CL | Batch | -0.013 | 0.011 | -1.183 | -0.529 | 1 |
| 3 | CL | Naive | -13 | 0.011 | -0.091 | -0.041 | 1 |
| 3 | Batch | Naive | 0.012 | 0.011 | 1.092 | 0.488 | 1 |
| 4 | CL | Batch | -0.002 | 0.011 | -0.182 | -0.081 | 1 |
| 4 | CL | Naive | 0.019 | 0.011 | 1.729 | 0.773 | 1 |
| 4 | Batch | Naive | 0.021 | 0.011 | 1.912 | 0.855 | 1 |
| 5 | CL | Batch | 0.019 | 0.011 | 1.729 | 0.773 | 1 |
| 5 | CL | Naive | 0.045 | 0.011 | 4.096 | 1.832 | 0.013 |
| 5 | Batch | Naive | 0.026 | 0.011 | 2.367 | 1.058 | 1 |
| 6 | CL | Batch | 0.012 | 0.011 | 1.092 | 0.488 | 1 |
| 6 | CL | Naive | 0.072 | 0.011 | 6.554 | 2.931 | < .001 |
| 6 | Batch | Naive | 0.06 | 0.011 | 5.462 | 2.442 | < .001 |

| | | | | | | | |
|---|---|---|---|---|---|---|---|
| 7 | CL | Batch | 0.022 | 0.011 | 2.003 | 0.896 | 1 |
| 7 | CL | Naive | 0.073 | 0.011 | 6.645 | 2.972 | < .001 |
| 7 | Batch | Naive | 0.051 | 0.011 | 4.642 | 2.076 | 0.002 |
| 8 | CL | Batch | 0.021 | 0.011 | 1.912 | 0.855 | 1 |
| 8 | CL | Naive | 0.077 | 0.011 | 7.009 | 3.134 | < .001 |
| 8 | Batch | Naive | 0.056 | 0.011 | 5.097 | 2.28 | < .001 |
| 9 | CL | Batch | 0.024 | 0.011 | 2.185 | 0.977 | 1 |
| 9 | CL | Naive | 0.101 | 0.011 | 9.194 | 4.111 | < .001 |
| 9 | Batch | Naive | 0.077 | 0.011 | 7.009 | 3.134 | < .001 |
| 10 | CL | Batch | 0.026 | 0.011 | 2.367 | 1.058 | 1 |
| 10 | CL | Naive | 0.107 | 0.011 | 9.74 | 4.356 | < .001 |
| 10 | Batch | Naive | 0.081 | 0.011 | 7.373 | 3.297 | < .001 |

Supplementary Table 4: Post hoc comparisons of the three training modes in each step in protocol 2 on the LFW benchmark. CL: CL + Replay, Batch, Naïve: CL Naïve

| # trained IDs | Training mode | Training mode | Mean Difference | SE | t | Cohen's d | $p_{tukey}$ |
|---|---|---|---|---|---|---|---|
| 3 | CL | Batch | -0.011 | 0.01 | -1.142 | -0.511 | 0.999 |
| 3 | CL | Naive | -0.01 | 0.01 | -1.038 | -0.464 | 1 |
| 3 | Batch | Naive | 0.0001 | 0.01 | 0.104 | 0.046 | 1 |
| 4 | CL | Batch | -0.017 | 0.01 | -1.764 | -0.789 | 0.945 |
| 4 | CL | Naive | -0.007 | 0.01 | -0.726 | -0.325 | 1 |
| 4 | Batch | Naive | 0.01 | 0.01 | 1.038 | 0.464 | 1 |
| 5 | CL | Batch | -0.027 | 0.01 | -2.802 | -1.253 | 0.336 |
| 5 | CL | Naive | -0.005 | 0.01 | -0.519 | -0.232 | 1 |
| 5 | Batch | Naive | 0.022 | 0.01 | 2.283 | 1.021 | 0.692 |
| 6 | CL | Batch | -0.004 | 0.01 | -0.415 | -0.186 | 1 |
| 6 | CL | Naive | 0.035 | 0.01 | 3.632 | 1.624 | 0.049 |
| 6 | Batch | Naive | 0.039 | 0.01 | 4.047 | 1.81 | 0.014 |
| 7 | CL | Batch | 0.016 | 0.01 | 1.66 | 0.743 | 0.968 |
| 7 | CL | Naive | 0.05 | 0.01 | 5.189 | 2.321 | < .001 |
| 7 | Batch | Naive | 0.034 | 0.01 | 3.528 | 1.578 | 0.065 |
| 8 | CL | Batch | 0.012 | 0.01 | 1.245 | 0.557 | 0.998 |
| 8 | CL | Naive | 0.05 | 0.01 | 5.189 | 2.321 | < .001 |
| 8 | Batch | Naive | 0.038 | 0.01 | 3.944 | 1.764 | 0.02 |

Supplementary Table 5: Comparison between computational models of familiar face recognition

| Topic | Kramer et al., 2018 | Blauch et al 2020 | Noyes et al., 2021 | Abudarham & Yovel |
|---|---|---|---|---|
| Face representation learning | Batch training | Batch training | Batch training | Continual learning |
| Learning a new familiar face | Retraining | Retraining | Retraining | No retraining. Insert embedding to memory |
| Unfamiliar face representation | PCA | DCNN penultimate layer (fc7) | DCNN penultimate layer (fc7) | DCNN output (arcface) |
| Familiar face representation | PCA+LDA | Probability layer | Averaging of fc7; Vector of SVM probabilities | DCNN output (arcface) |
| Familiar face recognition | Nearest neighbor | Output node | --- | Nearest neighbor |
| Familiar face matching | Recognize each face separately and match identities | Recognize each face separately using DCNN output node, and match identities. Or Prob layer similarity | Representation similarity (after SVM) | Recognize each face separately and match identities |
| Unfamiliar face matching | PCA similarity | Fc7 similarity | Fc7 similarity | DCNN output (arcface) similarity |
| Face space structure / how familiar faces are stored in memory | Single prototype, PCA + LDA of average image pixels | An output node for every familiar face | DCNN representations, use SVM to recognize familiar identity | Multiple prototypes, representation cluster centroid + std |
